# UHRF1, NSUN2, and NEIL1 were Detected as Clinical Biomarker Candidates in Prostate Adenocarcinoma with Potential Roles in Disease Pathogenesis

**DOI:** 10.1101/2021.10.23.465589

**Authors:** Buse Nur Kahyaoğulları, Turan Demircan

## Abstract

Prostate cancer (PCa) is the most commonly diagnosed cancer in men. Expanding evidence suggests a significant association between cancer progression and RNA modifications. However, our knowledge of the link between m5C and hm5C pathways with PCa is limited. Therefore, we aimed to explore the diagnostic and prognostic values of m5C and hm5C regulators in PCa. In this study, genetic alterations in m5C and hm5C regulators were identified using publicly available databases. Differentially expressed genes in these pathways between tumor and nontumor samples, correlation among m5C and hm5C pathway members, and prognostic value of the regulators were evaluated. Furthermore, enrichment of gene ontology (GO) terms and KEGG pathways was carried out. Obtained results unveiled the mRNA level differences as the key genetic alterations for m5C and hm5C regulators between tumor and nontumor samples. *UHRF1, TET3,* and *NEIL1* were significantly upregulated in tumor samples compared to nontumor ones, whereas *EGR1* was significantly downregulated. *UHRF1, DNMT1, NSUN2, NSUN4, C1orf77, C3orf37, WDR77, NEIL1*, and *TDG* genes were identified as candidate prognostic markers of overall survival. The upregulated genes in patient samples with genetic alterations in m5C and hm5C pathways enriched cell cycle-related processes. In summary, our findings suggest that the m5C and hm5C regulators might play a role in PRAD development by activation of proliferation, and the *UHRF1, NSUN2*, and *NEIL1* genes have the potential to be utilized as clinical biomarkers.

## Introduction

Prostate cancer (PCa) is the second most diagnosed cancer (13.5%) in men after lung cancer (14.5%), with a 6.7% death rate[1]. Incidence and mortality of prostate cancer increase with age [2, 3]. Biopsy, digital rectal examination, transrectal ultrasound, and prostate-specific antigen test are used to diagnose PCa [4]. Histological imaging is mainly used in the staging of PCa, and the Gleason scoring system based on the morphological structure of prostate cancer is commonly adopted [5]. In this scoring method, a Gleason score ≤6 represents a good prognosis (grade 1), seven is used for grades 2 and 3, eight refers to grade 4, and finally, nine and ten portrey the worst prognosis with grade 5 [6]. Current treatment options for PCa are androgen hormone deprivation therapy (ADT), surgical removal of the tumor, and radiotherapy [7]. Androgen hormone has a vital role in PCa progression, and therefore several treatments to target androgen hormone have been developed [8]. ADT has a prevalent use to treat PCa [9], and although patients initially respond positively to ADT, almost all patients eventually develop castrate-resistant prostate cancer [10]. Hence, identification of new potential candidate genes to cure the PCa is required.

Epitranscriptomics, employing next-generation sequencing to decode the RNA modifications, has recently become one of the emerging fields for researchers [11]. By now, more than 170 types of RNA modifications have been described [12], and among them, N6-methyladenosine (m6A), N1-methyladenosine (m1A), 5-methylcytosine (m5C), 5-hydroxymethylcytosine (hm5C), pseudouridine (Ψ), inosine (I), and uridine (U) are commonly detected mRNA modifications [13]. Due to RNA modifications, mRNA stability, export, localization, and translation rate are affected [13]. Proteins that regulate the dynamic and reversible RNA modifications are classified into three groups. Enzymes that add methyl groups are called ‘writers’, binding proteins that recognize and bind the methylated residues are grouped as ‘readers’, and the proteins which catalyze the demethylation reaction are known as ‘erasers’ [14].

m5C was first identified in DNA as one of the epigenetic regulation mechanisms [15]. Recently, m5C modification was also determined in various RNA species, including rRNA, tRNA, non-coding RNA, and mRNA, with potential effects on RNA export, ribosome assembly, translation, and RNA stability [16]. m5C modification is abundantly found in coding and non-coding regions along a transcript, suggesting that m5C modification may play a significant role in translation rate and the post-transcriptional regulation of the mRNA,[16]. m5C modification is provided by the writer enzymes member of DNA methyltransferases (DNMTs), and NOP2/SUN RNA methyltransferase family (NSUN), particularly by the DNMT2 and NSUN2 [17, 18]. Localization of DNMT2 as a shuttling protein in the nucleus and cytosol during the cell cycle indicates that this methyltransferase actively modifies nucleic acids in both cellular compartments, which has been validated experimentally for a variety of RNA types besides the DNA [17, 19]. Moreover, previous reports demonstrated that the NSUN family could target mRNA, rRNA, mitochondrial DNA, and non-coding RNAs, apart from tRNAs, to add a methyl group to the cytosine base [20]. Aly/REF Export Factor (ALYREF), a reader protein in the m5C pathway, takes a role in exporting mRNAs modified primarily by NSUN2 [21]. Based on the accumulated evidence, the ten-eleven translocation (TET) protein family is considered the eraser of the m5C pathway by catalyzing the oxidation reaction on m5C in DNA and RNA to form a hm5C group [22, 23]. On the contrary, TET enzymes are classified as ‘writers’ in the hm5C pathway. The formation of hm5C in mammalian RNAs is evident in the previous studies [23]. Although the mechanism is not fully understood yet, translation efficiency is increased for the mRNAs modified with hm5C [24].

RNA modifications affect many cellular processes and play a role in various human cancers [25]. The poor prognosis of bladder cancer due to the 5mC methylation of the *HDGF* gene is associated with the activity of NSUN2 writer protein [26]. In hepatocellular carcinoma (HCC), the detected copy number variation (CNV) of m5C regulators and their expression levels have been correlated with the progression of the disease [27]. In addition, 5mC modifications in HCC, catalyzed by NSUN4, and recognized by the reader ALYREF protein, resulted in dysregulation of cell cycle and mitosis and therefore accounted for HCC development [27]. In glioblastoma multiforme (GBM), 5mC modification on miRNAs decreases the binding efficiency of miRNAs to target mRNAs, and hence methylation of miRNAs increases the gene expression [28]. m5C modification in miRNA-181a-5p exhibited a tumor-growth effect and has been related to a poor prognosis of GBM [28]. In ovarian cancer, differential expression of m5C regulators was detected in subtypes of cancer, and they are used to characterize the ovarian cancer subtypes [29]. Increased hm5C modification by TET enzymes has been demonstrated to prevent melanoma development [30]. In another study, the hm5C level decreased significantly in thyroid cancer patients with TERT promoter mutation [31].

The current study examined diagnostic and prognostic values of m5C and hm5C RNA regulators in PCa using TCGA pan-cancer datasets. An in-depth bioinformatics analysis was employed to unveil the link between genetic alteration in RNA regulators and PCa progression. Expression level differences between tumor and non-tumor samples were investigated, and a list of differentially expressed genes (DEGs) was identified. Several genes whose altered expression levels were associated with poor PCa progression, were detected as potential prognostic biomarkers. We believe that the findings of this study would be useful to understand the roles of RNA modification regulators in PCa development, and some of these could serve as valuable biomarkers of PCa.

## Materials and Methods

### TCGA Pan-cancer PRAD Dataset

The expression and clinical data for 550 samples (498 tumor and 52 non-tumor samples) from 498 PRAD patients were extracted from the TCGA database [32] by using the RTCGA package [33] in the R environment.

### Genetic alterations for m5C and hm5C regulators

The cBioportal database [34] was used to identify genetic alterations of the m5C and hm5C regulators in PRAD patients. cBioportal is an open-access website containing multidimensional cancer genomic data and aims to provide researchers with access to various molecular profiles and clinical data. Frequency and type of genetic alterations for m5C and hm5C regulators were profiled on an oncoprint graph using the PRAD Pancancer dataset, and then we selected seventeen studies (Figure 1b, d) for analyzing alteration frequency.

**Figure 1.**
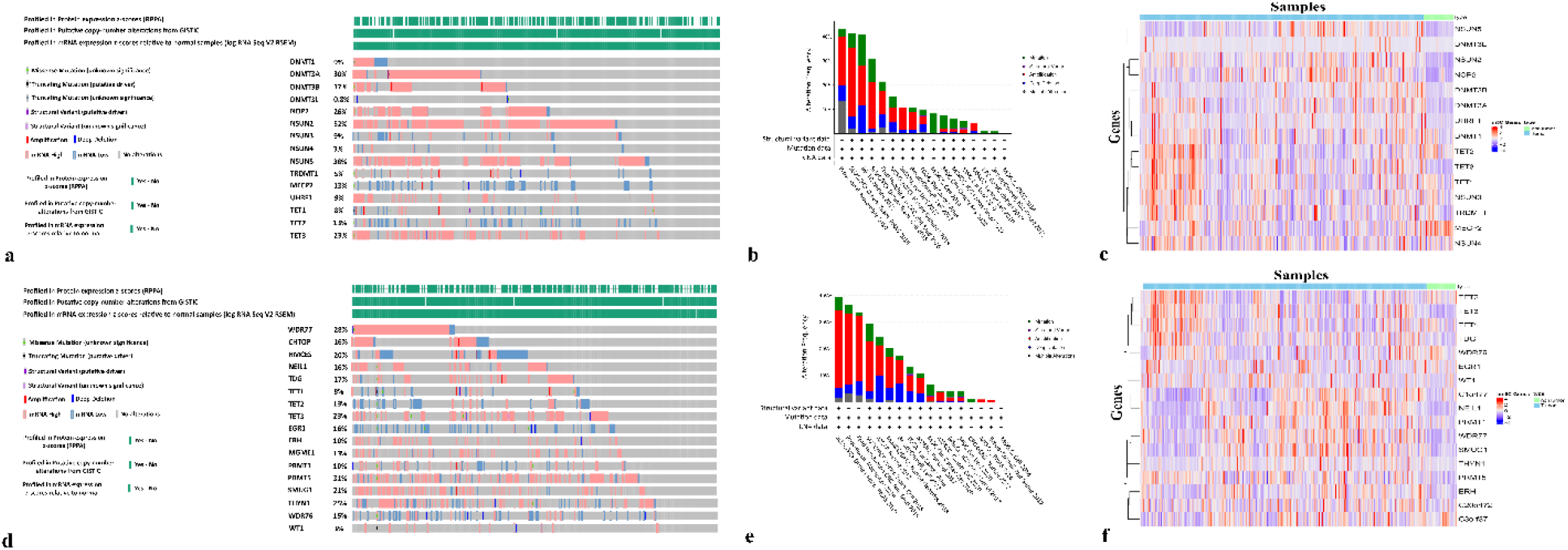
Genomic alterations of m5C and hm5C regulators in TCGA-PRAD cohort (494 samples). (a, d) Oncoprints of m5C and hm5C regulators for 15 and 17 genes in PRAD, respectively. (b, e) Cancer type summary for m5C and hm5C regulators in seventeen studies. (c, e) Expression levels of regulators in tumor and non-tumor samples of PRAD patients.

### Generated Heatmaps for m5C and hm5C Genes

To evaluate the expression levels of the m5C and hm5C regulators in prostate cancer, RNAseq data was extracted from the TCGA-PRAD cohort, and heatmaps were generated using ‘ComplexHeatmap’ package [35] in R language to display the expressions of m5C and hm5C regulators according to patients.

### Principle Component Analysis and Expression Level Differences of m5C and hm5C Regulators

Principal component analysis (PCA) was performed using the ‘ggfortify’ package of R language to assess the discrimination capacity of m5C and hm5C regulators’ expression levels between tumor and non-tumor samples. The same package was employed to visualize the results.

To test whether the expression levels of analyzed genes are significantly different in m5C and hm5C pathways between compared groups, the Wilcoxon test in ‘rstatix’ package was adopted. ‘ggplot’ package was implemented to display the results with significance levels on boxplots.

‘Limma’ package [36] in R language was exploited to identify the differentially expressed m5C and hm5C RNA methylation regulators between tumor and non-tumor samples. The cut-off was set as |FC| > 2 and p-value <0.05. The Differentially expressed genes (DEGs) in m5C and hm5C pathways were listed on separate tables.

### Correlation Analysis

Correlation maps and heatmaps were generated to explore the expression trend of the genes among tumor samples for m5C and hm5C pathways. Furthermore, a similar analysis was conducted using the unified list of the genes gathered from these two pathways. ‘Hmisc’ package was utilized for correlation analysis, and ‘corrplot’ package was used along with the base functions in the R language for visualization of the results. Correlation coefficient values were obtained between −1 to 1. The values close to −1 or 1 for the genes were considered as a strong correlation, whereas those close to 0 indicated a weak correlation. A positive correlation was shown with a value between 0 and 1, and −1 to 0 values signified the negative correlation.

### Kaplan-Meier overall survival analysis

For generated overall survival graphs, the clinical data (PRAD.clinical) was extracted from the ‘RTCGA.clinical’ package in the R environment. Kaplan-Meier estimation method was applied to test the prognostic value of the m5C and hm5C genes regarding the overall survival (OS) of prostate cancer patients with low or high m5C-hm5C gene expression. To determine the optimal cut-off and calculate the P-value for each gene ‘survival’ package in the R environment was used, and the list of obtained p-values was shown on a table. Significant genes (p-value <0.05) for m5C or hm5C pathways were selected to draw the Kaplan-Meier survival curves by utilization of ‘survminer’ and ‘ggplot2’ R packages.

### ROC Curves

Receiver operating characteristic (ROC) curve analysis was conducted utilizing the ‘pROC’ package [37] in R language to evaluate whether the expression level of selected m5C and hm5C regulators (*UHRF1, NSUN2*, and *NEIL1*) was distinctive for tumor and nontumor samples. Area under curve (AUC) value was calculated for each gene to estimate the sensitivity and specificity of the performed analysis.

### Expression Levels of *UHRF1*, *NSUN2*, and *NEIL1* Genes in PRAD Patients with Different Gleason Score

Gleason classification is used for morphological grading for prostate adenocarcinoma. The association of clinical features with expression levels of the genes was assessed through the UALCAN online database [38]. Expression levels in tumor samples with different Gleason scores (from 6 to 10) were compared with expression levels in non-tumor samples. The results were illustrated on the graphs generated by the UALCAN database.

### Enrichment Analyses for Altered and Unaltered Groups

We classified the tumor samples into two groups for both m5C and hm5C pathways. Tumor samples with genetic alterations for at least one gene formed the altered group and the unaltered group consisted of tumor samples without mutations for the genes in m5C or hm5C pathways. A similar grouping was used for the genes (*UHRF1*, *NSUN2*, and *NEIL1*) considered as putative prognostic markers.

Gene Ontology (GO) analysis was carried out to get all ontological terms enriched by DEGs for altered and unaltered groups. Molecular function (MF), biological process (BP), and cellular component (CC) terms were identified for each group with ‘clusterProfiler’ package [39] in R. The same package was implemented to unveil the Kyoto Encyclopedia of Genes and Genomes (KEGG) pathways in altered and unaltered groups. The applied parameters for GO and KEGG were as following: p-value <0.01, q-value <0.2, padjustmethod; ‘BH’

Statistically significant terms and pathways were visualized on dotplot and barplot graphs using R2s ‘ggplot2’ and ‘enrichplot’ packages as performed elsewhere [40].

### The communality of Enriched GO Terms for Altered and Unaltered Groups

Unique and common GO terms among altered and unaltered groups were found via Jvenn tool [41], as did elsewhere [42]. For both m5c and hm5C pathways, separate venn diagrams for altered and unaltered groups were generated.

### The Human Protein Atlas Data

The human protein atlas database [43] was utilized to map the expression of the selected proteins in normal and tumor tissues by immunohistochemical staining (IHC). Representative images of m5C (*UHRF1* and *NSUN2*) and hm5C (*NEIL1*) proteins in tumor and normal tissues were retrieved and visualized. While the proteins were labeled with protein-specific antibodies, a blue-colored hematoxylin dye was used to create contrast.

## Results

### m5C and hm5C Regulators were Found as Frequently Mutated in Prostate Cancer

Genetic alterations of m5C and hm5C pathway genes in prostate cancer were visualized on an oncoprint map using the cBioportal database (Figure 1a, d). For the m5C pathway, the top five genes with the highest alteration frequency were *NSUN2* (52%), *NSUN5* (38%), *DNMT3A* (30%), *NOP2* (26%), and *TET3* (23%) (Figure 1a). Besides *TET3*, which is an intersected member of both RNA modification pathways, *PRMT5* (31%), *WDR77* (28%), *THYN1* (25%), and *SMUG1* (21%) were found as commonly altered genes among hm5C regulators (Figure 1d).

Alteration profiles at genetic levels for m5C and hm5C regulators were demonstrated on barplots generated by the cBioportal database (Figure 1b, e). For both of the pathways, amplification was the most frequently observed alteration type, followed by mutation and deep deletion alterations (Figure 1b, d). Furthermore, it was found that the majority of the m5C and hm5C genes were upregulated in most of the tumor samples compared to non-tumor ones in PRAD patients (Figure 1c, f). Lower expression of MECP2 and WDR76 was noteworthy (Figure 1c, f).

### *UHRF1, TET3, NEIL1,* and *EGR1* were Identified as DEGs Between Tumor and non-Tumor Samples

PCA analysis was implemented to examine whether the m5C or hm5C regulators have a remarkable power to discriminate the tumor and nontumor samples (Figure 2a, d). As can be seen on PCA plots for the m5C (Figure 2a) and hm5C (Figure 2d) regulators, although the expression level of these genes has a considerable discrimination rate for both pathways (for m5C:PC1 = 28.8%, PC2 = 18.6%; for hm5C: PC1 = 24.2%, PC2 = 15.1%), tumor and non-tumor samples were still intermingled.

**Figure 2.**
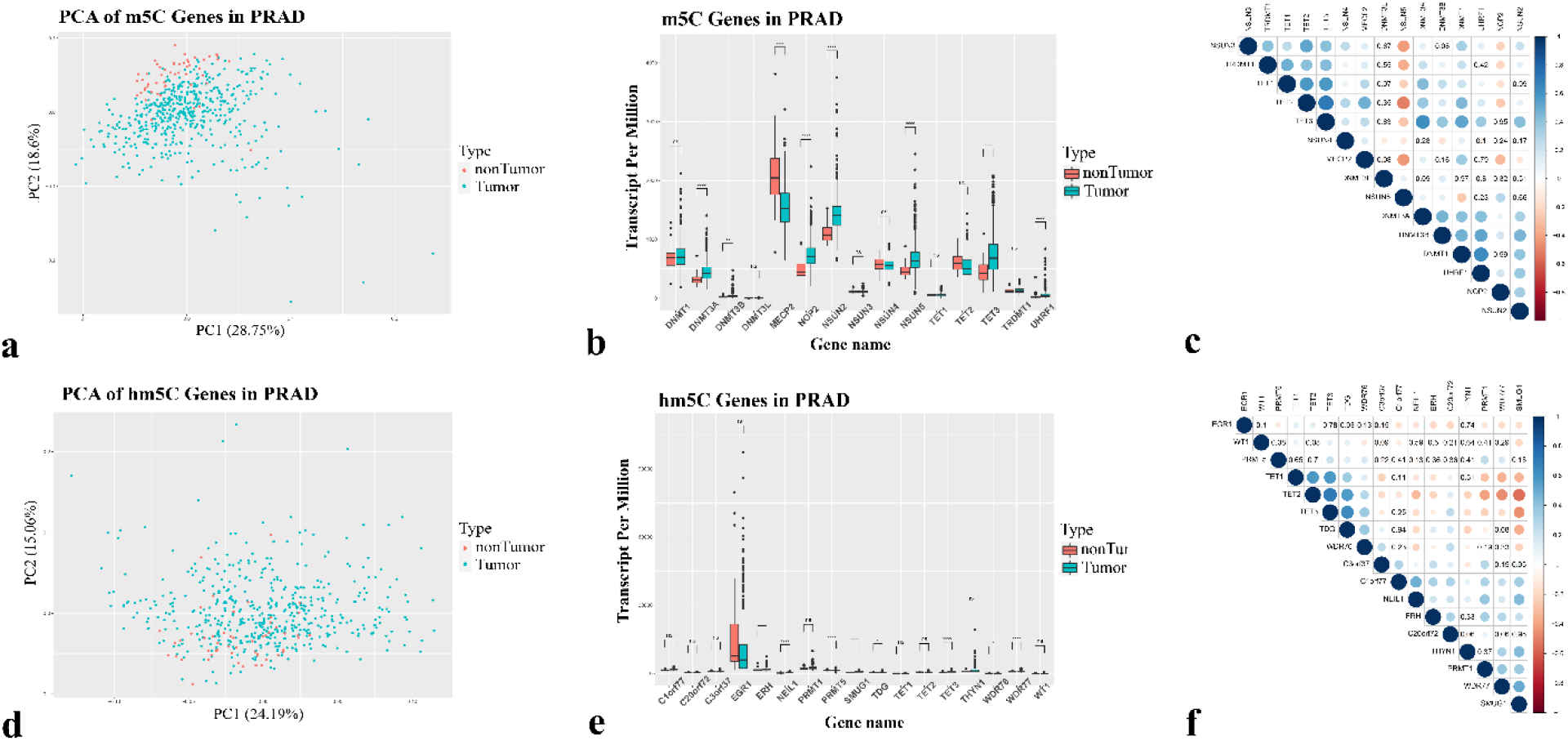
PCA, boxplots, and correlation maps for m5C-hm5C regulators. (a, d) PCA plots for m5C and hm5C genes, respectively. (b, e) Boxplots exhibit the expression levels of m5C and hm5C regulators between tumor and non-tumor samples. (c, f) Correlation of m5C and hm5C genes. Red and blue colors symbolize high and low correlation, respectively.

Then we applied the Wilcoxon test to calculate the p-value of each gene in m5C and hm5C pathways for tumor and nontumor samples without a fold change cut-off. Obtained results for each gene with significance level were displayed on boxplots (Figure 2b, e). Out of 5mC regulators, *DNMT3A, DNMT3B, NOP2, NSUN2, NSUN5, TET3,* and *UHRF1* expression levels were significantly higher in tumor samples compared to the nontumor group (Figure 2b). On the other hand, the expression level of the *MECP2* gene was detected as significantly lower in tumor samples (Figure 2b). The hm5C regulators *ERH, NEIL1, PRMT5, SMUG1, TDG, TET3, WDR76*, and *WDR77* were upregulated significantly in tumor samples (Figure 2e). Differential expression analysis between tumor and non-tumor samples using LIMMA resulted in one gene (*UHRF1*) in the m5C pathway and three genes (*TET3, NEIL1,* and *EGR*) in the hm5C pathway, all having an adjusted p-value < 0.01 and FC>1.5 (Supplementary Table 1). *UHRF1, TET3*, and *NEIL1* were significantly upregulated in tumor samples, and *EGR1* was identified as a significantly downregulated gene of RNA modification pathway genes in tumor samples compared to nontumor samples.

### Expression Levels of m5C and hm5C Regulatory Genes were Highly Correlated in PRAD Tumor Samples

To explore whether the genes in m5C or hm5C pathways were correlated ‘Spearman’ correlation test was applied to the analyzed genes, and the obtained results were presented in Figure 2. We observed a positive correlation trend for most of the genes in the m5C pathway (Figure 2c and Supplementary Table 2). Particularly, *TET2* and *TET3* (correlation coefficient: 0.71, p-value<0.001), *DNMT1* and *UHRF1* (correlation coefficient: 0.64, p-value<0.001), and *DNMT3A* and *TET3* (correlation coefficient: 0.62, p-value<0.001) were the most positively correlated gene pairs among the m5C regulators. In contrast, the top negatively correlated gene pairs were found as follows: *NSUN5* and *TET2* (correlation coefficient: −0.52, p-value<0.001), *NSUN5* and *MECP2* (correlation coefficient: −0.44, p-value<0.001), and *NSUN3* and *NSUN5* (correlation coefficient: −0.43, p-value<0.001).

On the other hand, for the hm5C pathway, the number of negatively and positively correlated gene pairs was close (Figure 2f and Supplementary Table 3). Aside from *TET2* and *TET3*, a positive correlation was manifested mainly in *TET3* and *TDG* (correlation coefficient: 0.64, p-value<0.001), *TET1* and *TET3* (correlation coefficient: 0.57, p-value<0.001), and *TET1* and *TET2* (correlation coefficient: 0.57, p-value<0.001) gene pairs. Contrary, the most negatively correlated gene pairs were detected as *TET2* and *SMUG1* (correlation coefficient: −0.55, p-value<0.001), *TET2* and *WDR77* (correlation coefficient: −0.46, p-value<0.001), and *TET3* and *SMUG1* (correlation coefficient: −0.45, p-value<0.001).

Correlation analysis was also conducted for the merged list of the genes in m5C and hm5C pathways (Supplementary Figure 1 and Supplementary Table 4.) to get insights into the potential cross-talk between these two pathways. The observed three clusters on the correlation heatmap indicated a strong positive correlation among the regulators of epitranscriptome pathways (Supplementary Figure 1). Relatively stronger positive correlation between the gene pairs *UHRF1* and *WDR76* (correlation coefficient: 0.78, p-value<0.001) and *DNMT1* and *WDR76* (correlation coefficient: 0.72, p-value<0.001) was an interesting finding. Notably, *MECP2* and *WDR77* (correlation coefficient: −0.51, p-value<0.001), *MECP2* and *SMUG1* (correlation coefficient: −0.47, p-value<0.001), and *NSUN3* and *SMUG1* (correlation coefficient: −0.46, p-value<0.001) gene pairs were revealed as negatively correlated genes in PRAD.

### *UHRF1, NSUN2,* and *NEIL1* were Identified as Potential Prognostic Biomarkers

The prognostic value of the m5C and hm5C regulators in PRAD was evaluated using the Kaplan-Meier estimation method (Supplementary Table 5). *UHRF1, DNMT1, NSUN2, NSUN4, C1orf77*, *C3orf37, WDR77, NEIL1,* and *TDG* genes were recognized as statistically significant (p-value<0.05) to predict the PRAD prognosis considering the overall survival data of patients (Figure 4a and Supplementary Figure 2). Low expression levels of *UHRF1, NSUN2, C1orf77, C3orf37, WDR77,* or *TDG* genes were associated with a better disease prognosis (Figure 3a and Supplementary Figure 2). However, low expression levels of *DNMT1, NSUN4*, or *NEIL1* genes were decreased the overall survival (Figure 3a and Supplementary Figure 2).

**Figure 3.**
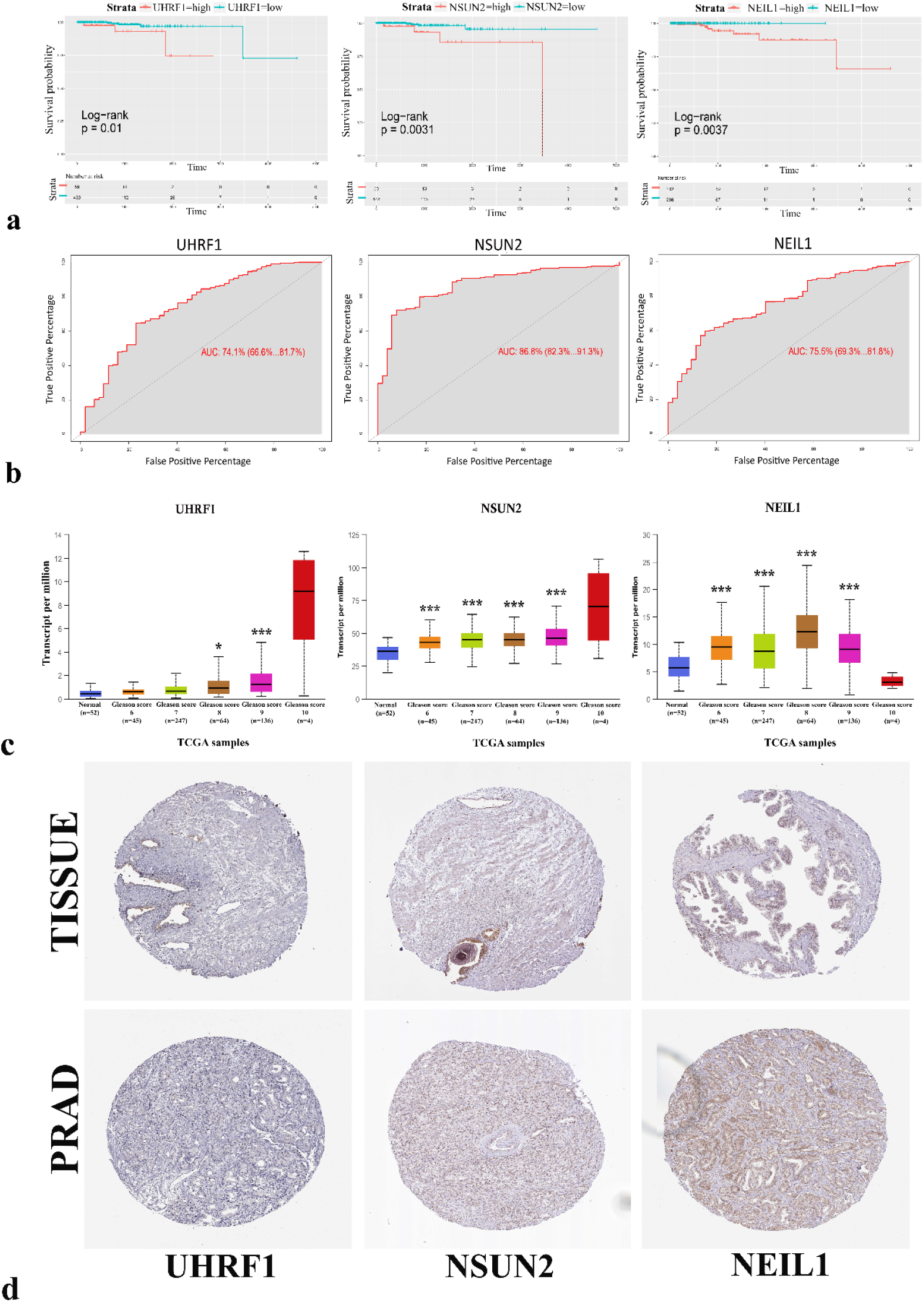
Represented pathways for altered and unaltered groups of m5C and hm5C genes. (a, c) Gene ontology (GO) analyses of m5C and hm5C altered groups, respectively. (b, d) Related pathways for m5C and hm5C unaltered groups on GO plots, respectively. BP; biological process, CC; cellular component, MF; molecular function.

The genes commonly identified in overall survival and LIMMA analyses (*UHRF1, NSUN2*, and *NEIL1*) were considered as candidate genes with potential prognostic value and roles in disease pathogenesis. Prediction scores of these genes were validated by plotting their ROC curves (Figure 3b). The area under the ROC curve (auROC) was calculated to estimate these genes’ sensitivity and specificity to distinguish the tumor and nontumor samples (Figure 3b). The auROC of the *UHRF1, NSUN2*, and *NEIL1* was calculated as %74.1, %86.8, and %75.5, respectively.

Gleason classification was performed in order to assess the correlation between mRNA expression levels of selected genes and clinical features of prostate cancer patients (Figure 3c). The expression level of the *UHRF1* gene was significantly higher in the samples with Gleason scores 8 and 9 compared to normal samples. Increased *NSUN2 and NEIL1* expression levels were also in accordance with Gleason grading scores. The expression levels of these genes were elevated in the samples with a higher Gleason score.

The Human Protein Atlas database was used to access IHC staining images of the *UHRF1, NSUN2*, and *NEIL1* genes to further examine how the expression level of m5C and hm5C regulators change in normal and tumor tissues. The selected three genes were more expressed in PRAD tissues (Figure 3d) than the nontumor samples. Increased IHC staining in the *UHRF1*, *NSUN2*, and *NEIL1* genes confirmed that the expression level of these three genes was increased in PRAD.

### Proliferation Related Biological Processes are Enriched in Altered m5C and hm5C Groups

To unveil the effect of m5C and hm5C regulators on disease pathogenesis, we focused on DE genes between altered and unaltered groups. The transcriptome data of patients with mutated genes in the 5mC pathway (altered group) was compared with the transcriptome data of patients without mutations in the 5mC pathway (unaltered group) using the cBioPortal database. The downloaded expression level list was filtered by setting p-value and fold-change cut-offs. Resultant genes were used to enrich the GO terms for altered and unaltered groups. The same approach was followed for hm5C pathway genes as well. GO enrichment analysis was separately carried out on the lists of altered and unaltered m5C and hm5C pathways groups.

A total of 145 biological processes (BP), 24 cellular components (CC), and 4 molecular functions (MF) were enriched by the 281 DEGs (Supplementary Table 6) in the altered group of the 5mC pathway. In contrast, the other 1810 DEGs in the unaltered group enriched a total of 607 BPs, 80 CCs, and 79 MFs (Supplementary Table 6). The 168 DEGs in the altered group of the hm5C pathway enriched a total of 93 BPs, 18 CCs, and 2 MFs (Supplementary Table 7), whereas the other 2204 DEGs in the unaltered group enriched a total of 893 BPs, 82 CCs, and 90 MFs (Supplementary Table 7).

Processes related to proliferation such as ‘chromosome segregation’, ‘nuclear division’, ‘sister chromatid segregation’, and ‘mitotic nuclear division’ were the top BPs enriched by the DEGs in the altered group of both m5C and hm5C pathways (Figure 4a, c), while ‘muscle contraction’, ‘extracellular matrix organization’, ‘extracellular structure organization’, and ‘muscle system process’ were the top BPs enriched by the DEGs in unaltered groups of both m5C and hm5C pathways (Figure 4b, d). Molecular functions such as ‘microtubule binding’ and ‘tubulin binding’ were the common MFs enriched by the DEGs in altered group of m5C and hm5C pathways (Figure 4a, c), however ‘passive transmembrane transporter activity’, ‘channel activity’, ‘ion-channel activity’, and ‘gated channel activity’ were among the common top MFs enriched by the DEGs in unaltered group of m5C and hm5C pathways (Figure 4a, c). On the other hand, ‘Chromosomal region’, ‘spindle’, ‘chromosome, centromeric region’, and ‘kinetochore’ were identified as the enriched top CCs in the altered group of m5C and hm5C pathways (Figure 4a, c), whereas ‘collagen-containing extracellular matrix’, ‘synaptic membrane’, and ‘transmembrane transporter complex’ were detected as the top CCs enriched by the DEGs in the unaltered group (Figure 4b, d).

**Figure 4.**
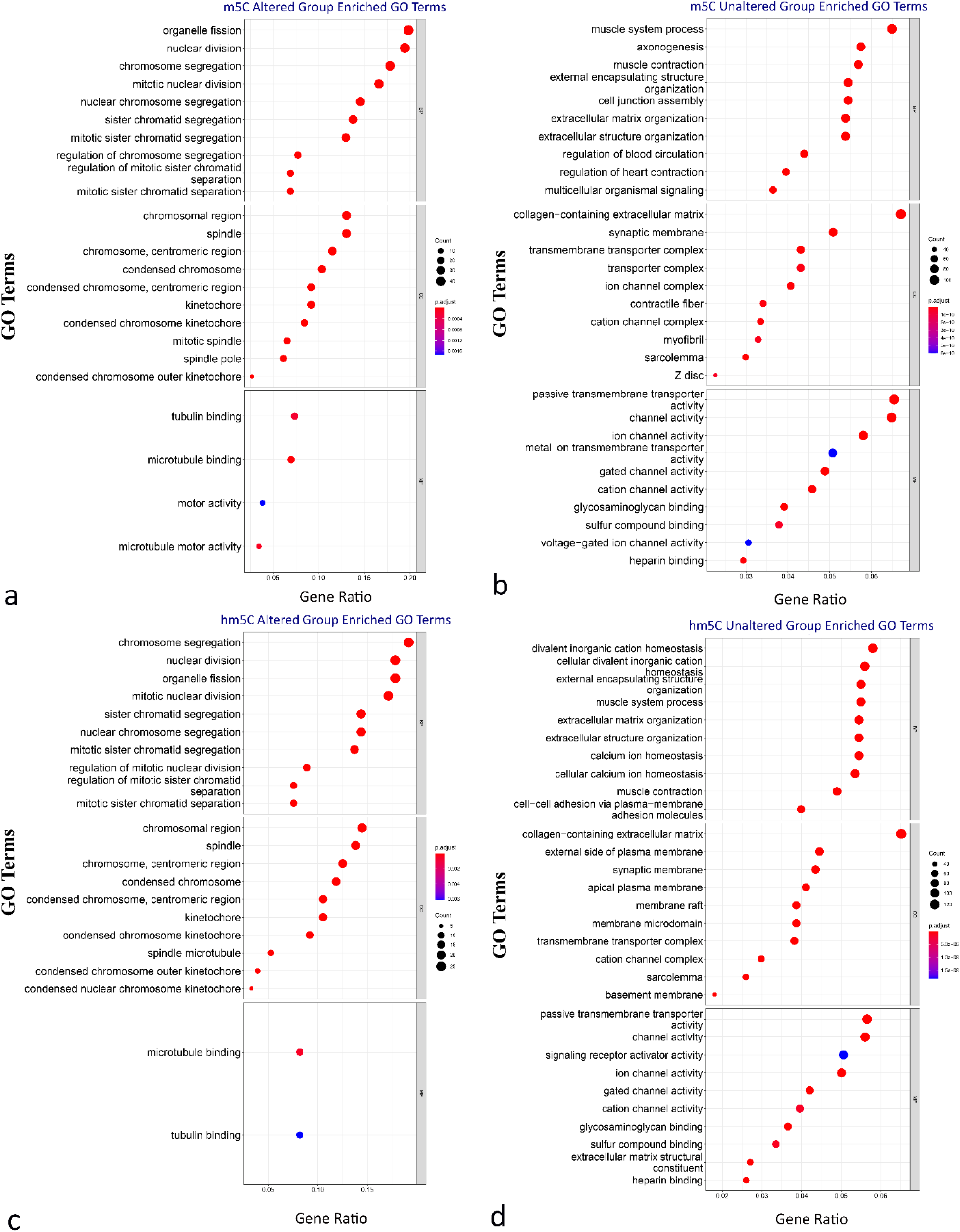
Potential biomarkers of PRAD (a, b) Overall survival and ROC curve analysis of selected genes. (c) Expression levels of *UHRF1, NSUN2,* and *NEIL1* in different Gleason scores. (d) Immunohistochemical staining images of *UHRF1, NSUN2,* and *NEIL1*

Enrichment of the ‘cell cycle’ and ‘oocyte meiosis’ KEGG pathways for both m5C and hm5C altered groups was remarkable (Supplementary Figure 3a, c). On the contrary, for the unaltered groups, ‘calcium signaling pathway’, ‘dilated cardiomyopathy’, and ‘hypertrophic cardiomyopathy’ pathways were the top enriched KEGG pathways (Supplementary Figure 3b, d).

To further dissect the putative roles of selected genes in PRAD pathology, we compared the gene expression profile of the PRAD patients classified according to the mutation status of the selected genes. As presented in Figure 5a, enriched terms for *UHRF1* or *NSUN2* altered groups were highly alike. ‘Organelle fission’, ‘nuclear division’, and ‘chromosome segregation’ were the top significant BPs, ‘mitotic spindle’ and ‘chromosomal region’ were the most significant enriched CCs, and ‘microtubule binding’, ‘motor activity’, and ‘ATPase activity’ were detected as the common MFs for *UHRF1* or *NSUN2* altered groups.

**Figure 5.**
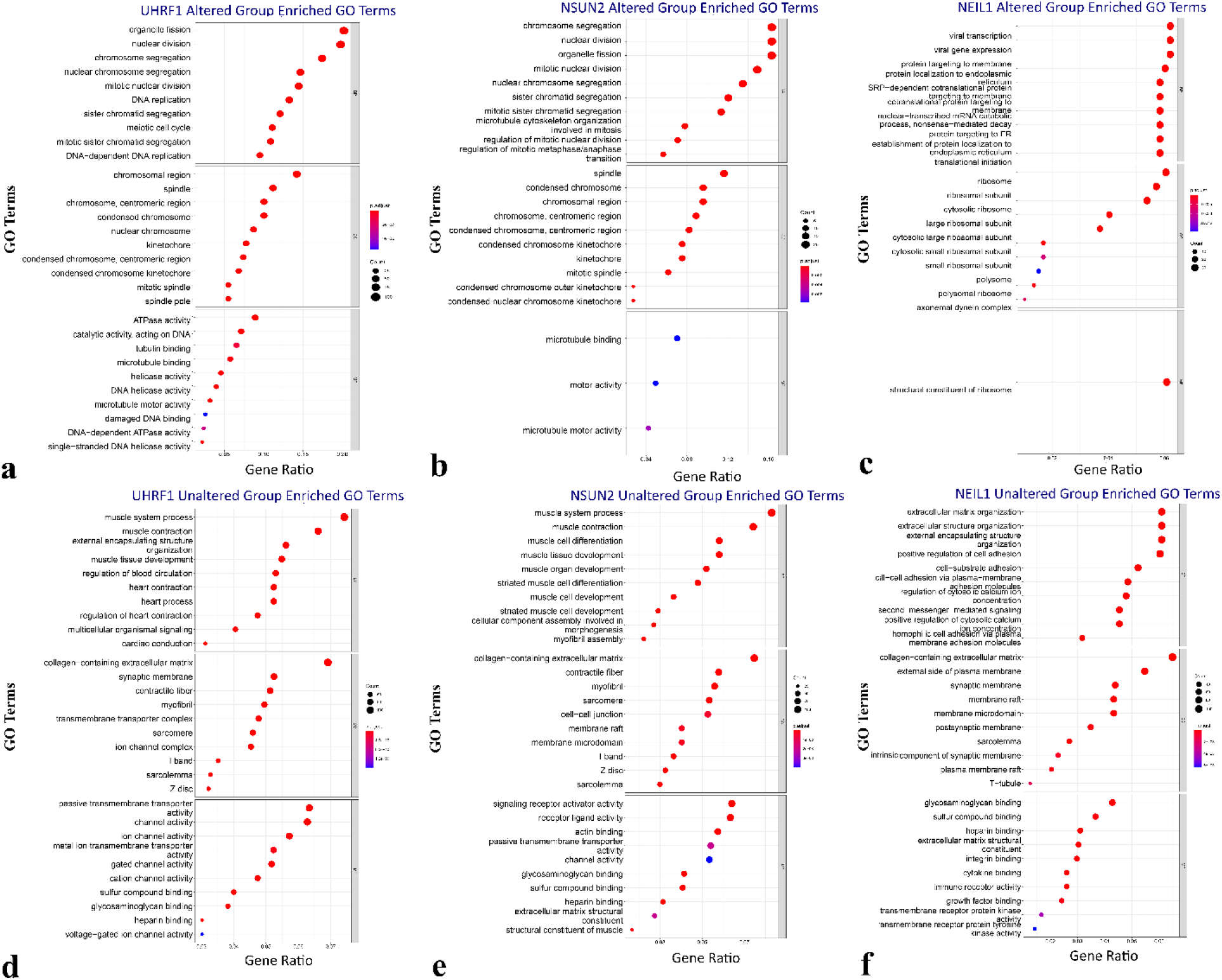
Enriched GO Terms for *UHRF1* altered (a) and unaltered (b), *NSUN2* altered (c) and unaltered (d), and *NEIL1* altered (e) and unaltered (f), groups. BP; biological process, CC; cellular component, MF; molecular function.

It is remarkable to observe that *NEIL1* altered group was diversified from the other two genes considering the enriched GO terms. For the *NEIL1* altered group, the most significant detected BPs were as follows: ‘viral transcription’, ‘viral gene expression’, and ‘protein targeting to membrane’ as BPs, ‘ribosome’, ‘ribosomal subunit’, and ‘polysome’ as CCs, and ‘structural constituent of ribosome’ as MF (Figure 5a).

Similar top GO terms were enriched in *UHRF1* or *NSUN2* unaltered groups, and the top-enriched GO terms were different in the *NEIL1* unaltered group (Figure 5b). BPs such as ‘muscle system process’, ‘muscle contraction’, and ‘muscle tissue development’, were the top BPs enriched by the upregulated genes in the unaltered group of *UHRF1* or *NSUN2*. Those upregulated genes in *NEIL1* unaltered group enriched ‘extracellular matrix organization’, ‘extracellular structure organization’, and ‘external encapsulating structure organization’ terms defined top enriched BPs.

Shared molecular functions such as ‘glycosaminoglycan binding’, ‘sulfur compound binding’, and ‘heparin binding’ were among the top MFs enriched by the genes in *UHRF1, NSUN2*, or *NEIL1* unaltered groups (Figure 5b). ‘Collagen-containing extracellular matrix’, ‘contractile fiber’, and ‘myofibril’ were among the top CCs enriched by the genes in *UHRF1* or *NSUN2* unaltered groups (Figure 5b), while ‘Collagen-containing extracellular matrix’, ‘external side of plasma membrane’, and ‘synaptic membrane’ were among the top CCs enriched by the genes in *NEIL1* unaltered group (Figure 5b).

KEGG results showed the ‘cell cycle’ was the common enriched pathway for *UHRF1* and *NSUN2* altered groups. ‘DNA replication’, ‘oocyte meiosis’, and ‘p53 signaling pathway’ were the examples of observed pathways only in the *UHRF1* altered group. In *NSUN2* altered group, the other pathway was ‘maturity onset diabetes of the young’ (Supplementary Figure 4a). In unaltered groups, *UHRF1* and *NSUN2* enriched pathways were alike. ‘calcium signaling pathway’, ‘dilated cardiomyopathy’, and ‘hypertrophic cardiomyopathy’ pathways were enriched in both unaltered groups (Supplementary Figure 4b). On the other hand, the profile of KEGG pathways enriched by the genes in altered and unaltered groups of *NEIL1* was remarkably different from *UHRF1* and *NSUN2* profiles (Supplementary Figure 4b). The only two KEGG pathways identified in the altered group were ‘ribosome’ and ‘coronavirus disease’, and the top three unaltered pathways were ‘cell adhesion molecules’, ‘hematopoietic cell linage’, and ‘malaria’.

### High Number of Shared Enriched BPs in m5C and hm5C Groups is Noteworthy

Shared and unique BP terms among the altered and unaltered groups were displayed on venn diagrams (Figure 6). m5C, *UHRF1*, and *NSUN2* altered groups had 63 shared BPs, whereas 167 BPs were found as unique to one of the altered groups (Figure 6a). For unaltered m5C, *UHRF1*, and *NSUN2* groups, 344 BPs were detected as common, and 367 BPs were identified as unique BPs (Figure 6b). Furthermore, the same analysis was applied for hm5C and *NEIL1* comparisons. Intriguingly, we found no shared BP between altered hm5C and *NEIL1* groups. On the other hand, 563 and 564 BPs were classified as shared or unique BPs between unaltered hm5C and *NEIL1* groups. The high number of shared BPs among unaltered groups was notable.

**Fig 6.**
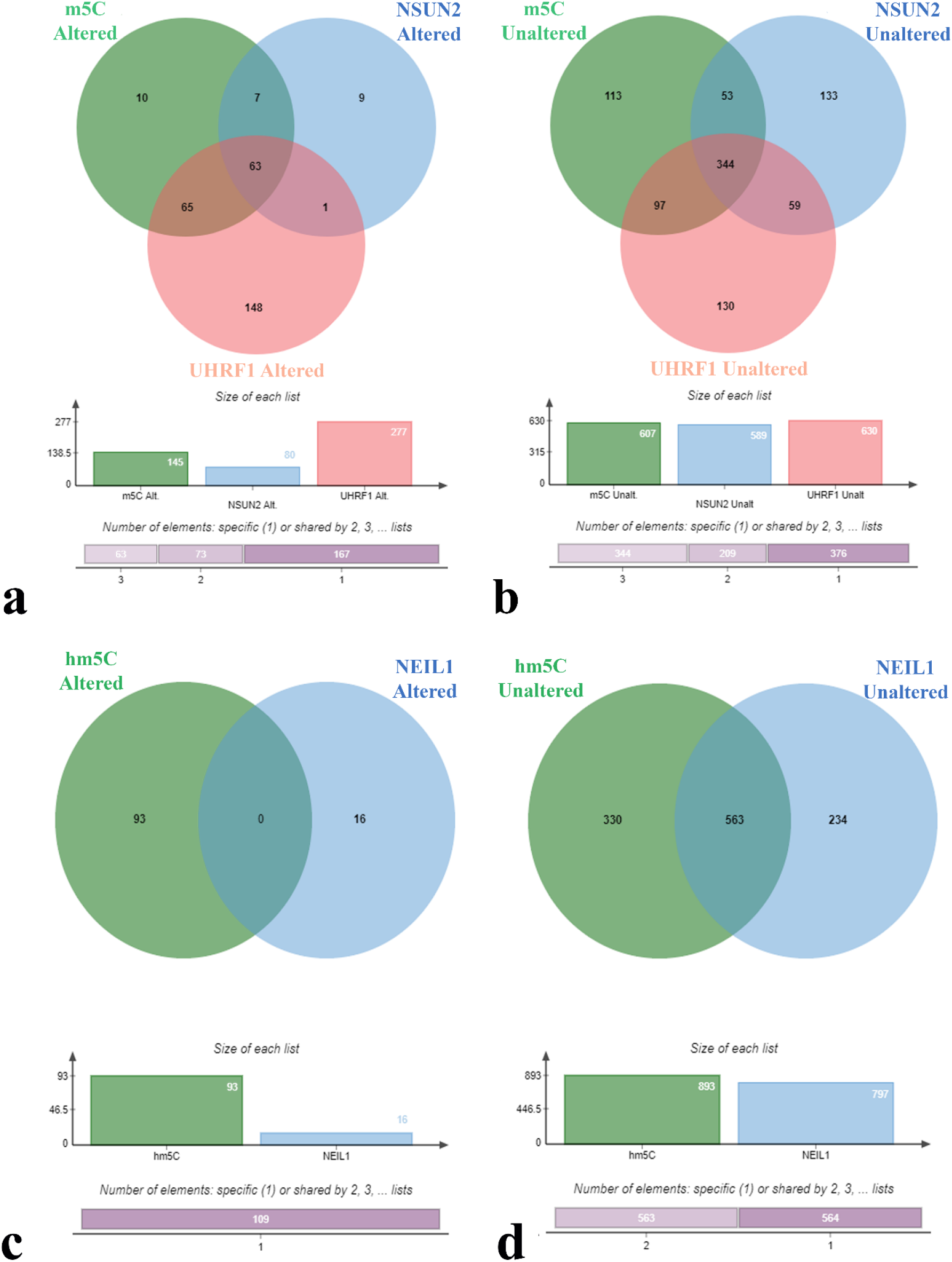
Commonly enriched and unique BPs among m5C, *UHRF1, NSUN2* altered (a) and unaltered (b), and between hm5C and *NEIL1* altered (c) and unaltered (d) groups.

## Discussion

PCa is one of the widely detected cancer types in man [44]. Although the death rate of PCa is not as high as some highly aggressive cancers due to its common diagnosis, studies aim to explore the pathology and prognosis of PCa has great importance in the cancer field. Recent studies have demonstrated that genetic alterations in epitranscriptomic regulators result in many diseases, including cancer [45]. Besides m6A, mutations in m5C and hm5C regulators have been also linked to cancer pathology. In this study, we examined diagnostic and prognostic values of hm5C and m5C regulators in PRAD.

mRNA level changes were the most frequently observed alteration type at the genetic level, whereas amplification was detected as the most common change at the DNA level. The uncovered high mutation rate and increased expression level of *NSUN2* in PRAD were consistent with previous cancer studies [46, 47]. The increased amplification and mutation levels of *NSUN2* in ovarian carcinoma, oesophageal carcinoma, bladder carcinoma (BLCA), cervical squamous cell carcinoma (CESC), lung squamous cell carcinoma (LUSC), and lung adenocarcinoma (LUAD) have been indicated in a pan-cancer study [47]. Moreover, in another study, a high *NSUN2* mRNA level in LUAD and LUSC compared to nontumor samples was found [46]. Overexpression of *NSUN2* in colorectal carcinoma (CRC) [48] and head and neck squamous cell carcinoma (HNSCC) [49] due to copy-number amplification was documented earlier.

Identification of *NSUN5* as another frequently altered regulator in PRAD is also well aligned with the literature. A pan-cancer study illustrated that *NSUN5* was overexpressed in various cancers, like glioblastoma multiforme, BLCA, PRAD, and breast invasive carcinoma (BIC) [50]. Furthermore, it has been documented that *NSUN5* promotes CRC development by enhancing cell proliferation [51], which is in line with another study underlining the confining effect of *NSUN5* deficiency on the proliferation and growth rate of mammalian cells [14]. We detected a high *NOP2*, also known as *NSUN1*, mRNA level in PRAD samples which ties well with previous findings. The amplification of *NOP2* and its overexpression is frequently found in renal cell carcinoma (RCC) [52]. Increased *NOP2* expression in diverse cancer types such as LUAD, LUSC, adrenocortical carcinoma, and ovarian serous cystadenocarcinoma was described in a reported pan-cancer study [53]. *NSUN2, NSUN5*, and *NOP2* are members of NOP2/SUN domain (NSUN) protein family, which act as DNA methyltransferase [17]. These enzymes participate in essential cellular functions such as cellular proliferation, cellular senescence, cellular migration, cellular differentiation, mRNA nuclear export, mRNA stabilization, tRNA stabilization, miRNA processing, and enhanced mRNA translation [46]. The significant roles of *NSUN2* and *NSUN5* in vital biological processes may explain the effect of dysregulation in these genes on PRAD development.

*DNMT3A*, a member of the DNA methyltransferase (DNMT) family, was identified as another m5C regulator with high mRNA expression level in PRAD samples. The increased mutation rate of *DNMT3A* in several cancers, including endometrial carcinoma (EC), acute myeloid leukemia (AML), LUAD, CESC, rectum adenocarcinoma, mesothelioma, was displayed in a previous pan-cancer study [54]. As a regulator of de novo methylation, *DNMT3A* functions in stem cell activity, proliferation, and differentiation [55]. Based on our analysis, the increased level of *DNMT3A* in PRAD samples provides evidence for the requirement of *DNMT3A* over-activation for cancer progression to alter the epigenetic methylation mark of cancer cells.

It was noteworthy to obtain high *TET3* and low *TET2* mRNA levels in PRAD samples. Ten-eleven translocation (TET) family members possess the catalytic activity to convert the m5C to hm5C. Therefore, TET proteins are not only considered as m5C erasers but are also accepted as writers of the hm5C pathway [14]. In many cancers, mutations in TET enzymes with different outcomes have been highlighted. Acting as a tumor-suppressor, *TET2* inhibits the BIC progression [56], and it is frequently mutated in hematopoietic malignancies [57]. On the other hand, higher *TET3* levels are associated with a poor prognosis of ovarian cancer [58] and BIC [59], whereas its downregulation promotes GBM progression [60]. Even though our results are generally consistent with older studies, further investigation of the *TET*s’ role in PRAD pathology is required. Moreover, high levels of *TET3* related proteins, *PRMT5* (a member of protein arginine methyltransferases family) and *WDR77* (androgen receptor cofactor p44), in PRAD patients are broadly in line with what has been reported in previous researches. The increased mutation rate and expression level of *PRMT5* was defined for many cancers, including uterine, ovarian, bladder, lung, glioma, and melanoma [61]. Another study revealed that inhibition of *WDR77* activity in CRC suppresses cancer cell growth [62]. The roles of *PRMT5* and *WDR77* in contribution to prostate growth, differentiation, and modulation by affecting the expression of androgen receptor-related genes [63], support the putative link between high expression of these genes with PRAD pathology. The observed high mRNA level of *SMUG1* in PRAD samples is in accordance with the previously reported findings in melanoma [64]. *SMUG1*, single-strand-selective monofunctional uracil-DNA glycosylase 1, is an eraser of the hm5C pathway [65]. As a base excision repair enzyme to catalyze the removal of the uracil and oxidized bases, *SMUG1* plays multiple roles to alter the RNA modification status and maintain genome stability [65]. Hence, its general cancer-suppressive roles contradict the high expression profile in PRAD samples which could be related to additional uncovered molecular functions of *SMUG1* that should be interrogated in future studies. Taken together, high genetic alteration rates of m5C and hm5C regulators in PRAD detected in this study are broadly consistent with previous findings.

The list of identified DEGs for m5C and hm5C pathways was noteworthy. Some of the readers, writers, or erasers exhibited different expression trends despite having similar activities. The observed opposite trend of the regulators with similar activity might be due to the expression under specific conditions or acting on different targets. Also, the expression trend of m5C and hm5C regulators may differ among various cancer types. For instance, *DNMT3A* is found as overexpressed in gastric adenocarcinoma [66], pancreatic adenocarcinoma (PDAC) [67], AML [68], gastroenteropancreatic cancer [69], HCC [70]. On the contrary, the loss of *DNMT3A* is responsible for a preleukemic phenotype on murine hematopoietic stem cells [71]. Downregulation of *DNMT3A* leads to tumor growth in BIC and CRC cell lines [72], and the lack of *DNMT3A* was related to tumor progression in LUAD [73]. Therefore, complementary to pathway scale examination, each gene in the RNA modification pathways should be evaluated individually.

The performed analysis of m5C and hm5C regulators based on gene expression levels in PRAD patients pinpointed a strong positive correlation among TET enzymes, which was also reported in earlier studies conducted in gastric cancer [74] and HNSCC [75]. Findings of positive correlation among DNMT enzymes, and between *TET3* and *DNMT3A* genes support the previous results in HNSCC [75] and non-small cell lung carcinoma [76]. Likewise, the detected negative correlation between *NSUN5* and *TET2*, and *NSUN5* and *TRDMT1* in this study were well aligned with the obtained data in pan-cancer [54] and RCC [77] studies. These results indicated that interactions among m5C and hm5C regulators might play crucial roles in PRAD. The significance of these correlations among the m5C and hm5C regulators is an open question.

Interestingly, enriched top BPs in the altered m5C and hm5C pathways groups were highly similar and related to cell division, such as chromosome segregation, nuclear division, and mitotic nuclear division. Affected proliferation-related pathways in patients with genetic alteration in m5C and hm5C imply the potential roles of these pathways in PRAD progression. Previous in-silico studies suggested that the cancer patients with a mutation in mRNA modification regulators upregulate the cell cycle-related pathways in gastrointestinal cancer [78] and BIC [42]. Experimental evidence demonstrated that an increased level of the m5C regulator *NSUN2* enhances the proliferation of gastric cancer cells [79]. Moreover, *PRMT5* plays an essential role in epithelial-mesenchymal transition in pancreatic cancer by regulating the EGFR and B-catenin pathways [80]. Knock-down of *PRMT5* in the pancreatic cancer cell line inhibits the growth of the cancer cells [25]. Since the hm5C pathway has not been extensively studied, we were not able to compare GO results for PRAD results with other cancer types due to the lack of reports on the enrichment of gene ontology terms considering the mutation status in hm5C regulators in different cancers.

Relation of the expression level of m5C and hm5C regulators and OS of PRAD patients was assessed to explore the prognostic value of these genes. Among the examined genes, *UHRF1, DNMT1, NSUN2, NSUN4, C1orf77, C3orf37, WDR77, NEIL1*, and *TDG* genes were detected as putative prognostic biomarkers. The genes common in DEG and OS lists (*UHRF1, NSUN2*, and *NEIL1*) were considered as strong candidates, which is confirmed by ROC curve analysis. The association of expression levels of these genes with overall survival as prognostic markers has also been demonstrated previously in multiple cancers. For instance, high levels of *UHRF1* resulted in lower overall survival in melanoma [81], HCC [82], and LUAD [82]. *NSUN2* high expression was also related to poor survival rates in hHNSCC [83], gastric cancer [79]. High *NEIL1* expression is correlated with low overall survival in gastric cancer [84]. Consistency of our findings with earlier reports prompted us to check whether *UHRF1, NSUN2*, and *NEIL1* expression levels of these genes are compatible with Gleason classifications. According to the obtained scores, we can conclude that the elevated expression levels of these genes are associated with clinically poor prognosis except the grade 10. The low sample number of Grade 10 is a limiting factor for a statistical evaluation of the relationship between expression level and Gleason score. High expression in PRAD was further validated by the comparison of protein expression profiles of *UHRF1, NSUN2*, and *NEIL1* in tumor and nontumor tissues using the Human Protein database. Taken together, we believe that *UHRF1, NSUN2*, and *NEIL1* can be considered as promising diagnostic and prognostic biomarkers of PRAD.

Therefore, we performed GO and KEGG analyses for altered and unaltered groups of *UHRF1, NSUN2*, and *NEIL1* genes to provide new clues on the impact of RNA modification pathways in PRAD development. BPs enriched by the upregulated genes in *UHRF1* and *NSUN2* mutated patients were greatly alike and considerably resembled the BPs obtained by the altered group of m5C. Most of the common altered BPs among m5C, *UHRF1*, and *NSUN2*, such as chromosome segregation, nuclear division, meiotic cell cycle processes, were related to cell cycle and proliferation. *UHRF1* has roles in *DNMT1* degradation by functioning as an E3-ubiquitin-ligase [85]. Therefore, for many cancer types, global DNA hypomethylation occurs through the upregulation of *UHRF1* [86]. It has been reported that *UHRF1* participates in the development and progression of many cancers, such as gastric cancer [87], non-small lung carcinoma [88], and EC [89]. Moreover, its high expression is associated with a poor prognosis in bladder cancer [90], pancreatic cancer [89], and HCC [87]. Likewise, the pivotal roles of *UHRF1* in PRAD development have been experimentally corroborated [88, 91]. A similar association with cancer progression has been also described for *NSUN2* overexpression. High expression of *NSUN2* was linked to the development of several cancers such as esophageal squamous cell carcinoma (ESCC) [92], gastric cancer [79], and hypopharyngeal squamous cell carcinoma and [93]. Since our results indicated the high mRNA expression level as the most commonly detected alteration of *UHRF1* and *NSUN2* genes in PRAD patients, the obtained cell cycle-related BPs for the altered group of *UHRF1* and *NSUN2* strongly favors the findings of previously conducted studies. On the other hand, surprisingly, no common BPs were found between the *NEIL1* and hm5C altered groups. Diverse BPs enriched in altered *NEIL1* gene, and hm5C regulators groups imply the limited contribution of *NEIL1* to enrich the cell cycle-related pathways. Considering the *NEIL1* roles in the removal of carboxyl- and formylcytosine residues to alter the epigenetic profile [94], further functional studies should be addressed to describe its association with PRAD pathology.

In conclusion, our study comprehensively analyzes the m5C and hm5C regulators in PRAD development and progression. Our findings revealed that regulators in m5C and hm5C pathways are frequently mutated, and a high mRNA expression level was detected as a general mark in tumor samples. *UHRF1, NSUN2*, and *NEIL1* genes appeared as candidate biomarkers to use in PRAD. However, to further support the in-silico results of the present study, future in-vitro and in-vivo researches are required to elucidate the relationship between PRAD development and RNA modification mechanisms.

## Author Contributions

### Buse Nur Kahyaoğulları

Methodology, Formal Analysis, Investigation, Writing-Original Draft, Visualization.

### Turan Demircan

Conceptualization, Methodology, Formal Analysis, Investigation, Writing-Original Draft, Writing-Reviewing&Editing.

